# Perforin-2 is dispensable for host defense against *Aspergillus fumigatus* and *Candida albicans*

**DOI:** 10.1101/2024.09.23.614582

**Authors:** Mariano A. Aufiero, Li-Yin Hung, De’Broski R. Herbert, Tobias M. Hohl

**Affiliations:** Louis V. Gerstner Jr. Graduate School of Biomedical Sciences, Sloan Kettering Institute, Memorial Sloan Kettering Cancer Center, New York, NY, USA; Department of Pathobiology, School of Veterinary Medicine, University of Pennsylvania, Philadelphia, PA 19104, USA; Infectious Disease Service, Memorial Sloan Kettering Cancer Center, New York, NY, USA; Human Oncology and Pathogenesis Program, Memorial Sloan Kettering Cancer Center, New York, NY, USA

## Abstract

Myeloid phagocytes are essential for antifungal immunity against pulmonary *Aspergillus fumigatus* and systemic *Candida albicans* infections. However, the molecular mechanisms underlying fungal clearance by phagocytes remain incompletely understood. In this study, we investigated the role of perforin-2 (*Mpeg1*) in antifungal immunity. We found that *Mpeg1*^-/-^ mice generated on a mixed C57BL/6J-DBA/2 background exhibited enhanced survival, reduced lung fungal burden, and greater neutrophil fungal killing activity compared to wild-type C57BL/6J (B6) mice, suggesting that perforin-2 may impair antifungal immune responses. However, when we compared *Mpeg1*^-/-^ mice with co-housed *Mpeg*^+/+^ littermate controls, these differences were no longer observed, indicating that initial findings were likely influenced by differences in the murine genetic background or the microbiota composition. Furthermore, perforin-2 was dispensable for antifungal immunity during *C. albicans* bloodstream infection. These results suggest that perforin-2 is not essential for host defense against fungal infections in otherwise immune competent mice and highlight the importance of generating co-housed littermate controls to minimize murine genetic and microbiota-related factors in studies of host defense mechanisms.

**IMPORTANCE:** *Aspergillus fumigatus* is the leading cause of invasive aspergillosis (IA), which is associated with significant mortality, particularly in immunocompromised patients such as those with acute leukemia or undergoing hematopoietic stem cell transplants, where death rates reach 40-50% despite standard care. Treatments for IA remain limited and resistance to antifungals is emerging, leading the World Health Organization to recently classify *A. fumigatus* as a critical priority fungal pathogen. A greater understanding of how the immune system clears *A. fumigatus* could lead to host-directed therapies that could complement our current armamentarium of antifungal drugs and improve patient outcomes. Our findings reveal that perforin-2 is not essential for antifungal immunity against *A. fumigatus* in otherwise immune-competent mice and underscore the necessity of using co-housed littermate controls to avoid confounding factors in immunological studies.

## INTRODUCTION

*Aspergillus fumigatus* is a saprophytic mold that is ubiquitous in the environment and is the most common cause of invasive aspergillosis (IA), a disease with significant mortality even with standard of care (1). Following inhalation, *A. fumigatus* conidia (spores) enter lung alveoli and germinate, forming tissue invasive hyphae and leading to respiratory failure (2). Phagocytic cells of the myeloid lineage are essential for anti-*Aspergillus* immunity (3–5). Myeloid phagocytes express an array of effector molecules to kill *A. fumigatus* conidia following internalization and to inhibit the growth of hyphae that cannot be phagocytosed.

In corneal *A. fumigatus* infections where hyphae predominate, immune cells inhibit hyphal growth by limiting essential metal nutrients. Neutrophil-produced calprotectin (*S100A9*) chelates zinc and manganese, inhibiting hyphal growth *in vitro* (6). Conversely, calprotectin is redundant for neutrophil killing of *A. fumigatus* conidia *in vitro* and in the lung. Thus, *S100A9*^*-/-*^ mice fail to control hyphal growth during corneal infection, but effectively clear lung infection (6). Neutrophil-secreted lactoferrin inhibits *A. fumigatus* growth via iron deprivation, though its role in killing internalized conidia remains unclear (7). Phagocytes also produce various hydrolases with microbicidal activity. Neutrophil elastase and cathepsin G are crucial for murine survival after intravenous (i.v.) *A. fumigatus* infection (8). Elastase- and cathepsin G-deficient mice show elevated kidney fungal burden, the primary organ affected following intravenous infection, suggesting that these molecules are important for fungal killing, though this conjecture has not been directly tested. Acidic mammalian chitinase (*Chia*) is essential for neutrophil inhibition of *A. fumigatus* hyphal growth *in vitro*, and *Chia*^*-/-*^ mice exhibit higher fungal burden during corneal infection (9). In the lung, *Chia*^*-/-*^ mice show a slightly lower fungal burden than control mice, suggesting that *Chia* may be dispensable or inhibitory for pulmonary immunity (10), and its role in killing internalized *A. fumigatus* conidia remains untested.

NADPH oxidase is an essential antifungal molecule that produces reactive oxygen species (ROS) to kill fungi. NADPH oxidase-deficient murine neutrophils are less effective at killing internalized *A. fumigatus* conidia in the lung (11), and neutrophils from patients with genetic NADPH oxidase deficiencies are similarly impaired in killing *A. fumigatus* hyphae (12). Beyond NADPH oxidase-generated ROS, phagocyte mitochondria also produce ROS after *A. fumigatus* internalization. Mitochondrial ROS enhances *A. fumigatus* conidia killing by alveolar macrophages, but not by neutrophils, indicating a cell type-specific role in fungal killing (13). These findings underscore the diverse and context-dependent strategies phagocytes employ to combat *A. fumigatus*. However, our understanding of how phagocytes kill *A. fumigatus* during infections remains incomplete.

While the most substantial evidence supports the critical role of NADPH oxidase, the conidial killing defect of NADPH oxidase-deficient neutrophils is incomplete (11) and humans with congenic defects in the NADPH oxidase complex, termed chronic granulomatous disease, have only a 40-55% lifetime risk of developing IA despite universal environmental exposure (14). This finding suggests that additional mechanisms are involved in mediating fungal killing. In pursuit of uncovering novel antifungal mechanisms, we focused on perforin-2 (*Mpeg1)*, a member of the Membrane Attack Complex, Perforin/Cholesterol-Dependent Cytolysin (MACPF/CDC) superfamily of pore-forming proteins. Perforin-2 is highly expressed by phagocytic cells and localizes to pathogen-containing phagosomes (15, 16). Bacterial cells incubated with wild-type (WT) macrophages exhibit pores on their cell membranes; these are absent in bacterial cells from perforin-2-knockout macrophages (15). While this suggests perforin-2 can form pores on bacterial membranes, direct evidence of bacterial cell damage or lysis by perforin-2 alone is lacking. The role of perforin-2 in antibacterial immunity remains controversial, with conflicting reports on the susceptibility of *Mpeg1*^*-/-*^ mice to various bacterial infections (15–17). Similarly, the contribution of perforin-2 to microbial killing by phagocytes is disputed, with some studies reporting defective killing in *Mpeg1*^*-/-*^ neutrophils and macrophages (15, 18), while others found no such defect (16). Beyond its potential antibacterial role, perforin-2 has been implicated in dendritic cell function, facilitating IL-33 release during helminth infection (19) and cross-presentation of exogenous antigens (20). To facilitate cross-presentation, perforin-2 associates with antigen-containing endosomes and undergoes proteolytic cleavage upon fusion with lysosomes, allowing antigen leakage into the cytosol without affecting endolysosomal pH or proteolytic capacity (20).

Given the important role of phagocytes and intra-phagosomal killing to anti-*Aspergillus* immunity, we hypothesized that perforin-2 may contribute to anti-*Aspergillus* immunity. In the present study, we sought to clarify the contribution of perforin-2 to myeloid phagocyte fungicidal activity, fungal clearance, and murine survival during pulmonary *A. fumigatus* infection.

## RESULTS

To test the idea that perforin-2 contributes to antifungal defense, we infected *Mpeg1*^*-/-*^ or C57BL/6J (B6) wild type mice with a 4-6 × 10^7^ *A. fumigatus CEA10* conidia intratracheally (i.t.) and monitored the mice for survival and assessed fungal burden at 24 hpi. We found that while 60% B6 control mice succumbed to infection with *A. fumigatus, Mpeg1*^*-/-*^ mice were resistant to infection (Fig. 1 A). Consistent with improved survival following infection, *Mpeg1*^*-/-*^ mice had a reduced lung fungal burden at 24 hpi (Fig. 1 B). Together, these results suggest that perforin-2 may be detrimental for murine survival and clearance of *A. fumigatus* following infection, which was unexpected given that *Mpeg1*^*-/-*^ mice were previously shown to be susceptible to bacterial infection (15).

**Figure 1.**
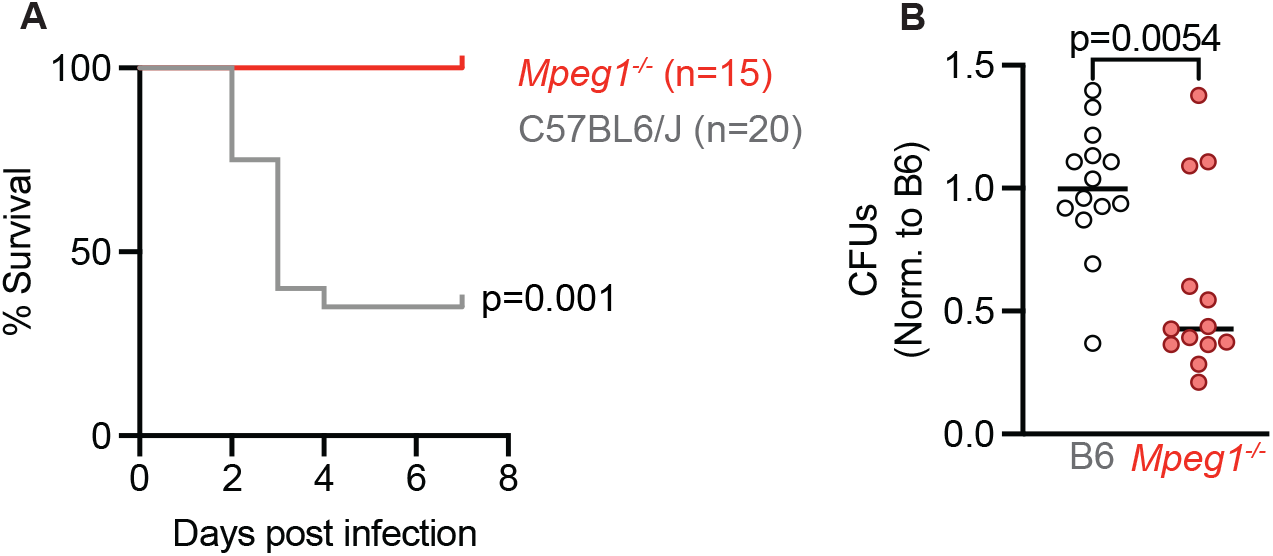
Mpeg1^-/-^ *mice have improved survival and fungal clearance following* A. fumigatus *lung infection compared to B6 mice*. (A) Survival of B6 and *Mpeg1*^*-/-*^ mice after infection with 4-6×10^7^ *A. fumigatus*. conidia. Significance calculated by log-rank (Mantel-Cox) test. (B) CFU from lungs of B6 and *Mpeg1*^*-/-*^ mice at 24 hpi with 3×10^7^ conidia. Each dot represents a mouse, and the bar indicates mean. Significance calculated by Mann-Whitney test. All data are pooled from 2 experiments.

To explore this phenotype further, we hypothesized that perforin-2 may be detrimental to murine survival by limiting the recruitment or antifungal activity of phagocytes in the lung. To test this hypothesis, we infected mice i.t. with Fluorescent *Aspergillus* Reporter (FLARE) conidia (11). FLARE conidia constitutively express red fluorescent protein (RFP) and are labeled with Alexa fluor 633 (AF633). Following uptake by phagocytes, fungal killing results in loss of RFP, while the AF633 is retained, allowing for the identification and quantification of phagocytes containing live or dead conidia. At 24 hpi, we isolated the lungs of FLARE infected mice, and quantified the total number of phagocytes as well as conidial uptake and conidial viability in these phagocytes. We did not see significant differences in the numbers of alveolar macrophages, neutrophils, or monocytes in *Mpeg1*^*-/-*^ mice compared to B6 control mice. However, we did see an increase in monocyte-derived dendritic cells (Mo-DCs) in *Mpeg1*^*-/-*^ mice (Fig. 2 A). We found that MoDCs from *Mpeg1*^*-/-*^ mice had greater fungal uptake compared to B6 control mice and inflammatory monocytes had a slight decrease in conidial uptake (Fig. 2 B and C). We also found that all lung phagocytes examined in *Mpeg1*^*-/-*^ mice had a decrease in conidial viability compared to B6 control mice (Fig. 2 B and D). These findings suggested that perforin-2 impaired the antifungal activity of phagocytes during *A. fumigatus* infection.

**Figure 2.**
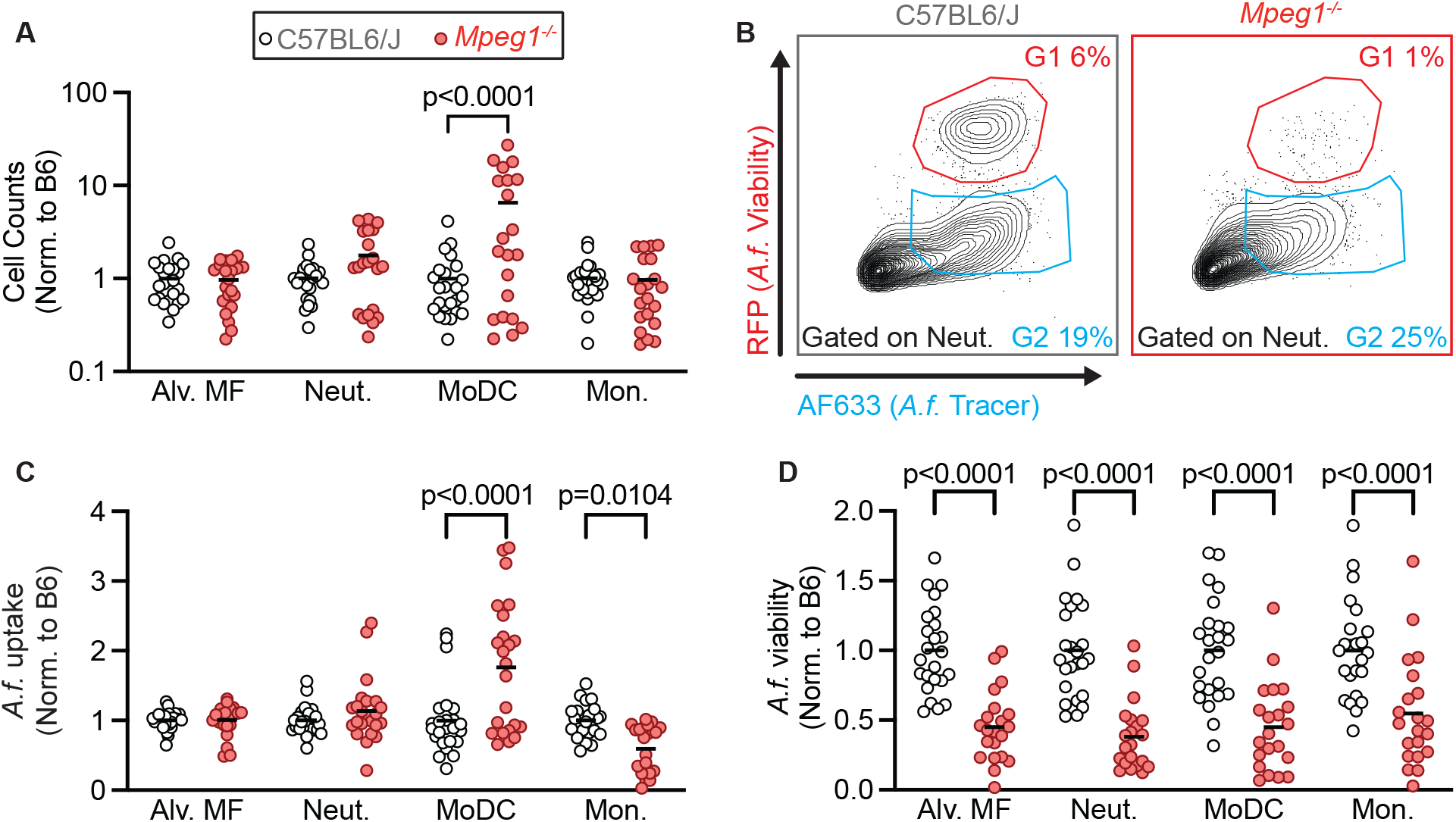
Mpeg1^-/-^ *phagocytes have superior antifungal activity relative to B6 mice*. (A) Cell counts in the lungs of B6 and *Mpeg1*^*-/-*^ mice of alveolar macrophages (Alv. MF), neutrophils (Neut.), MoDCs, and monocytes (Mon.) 24 hpi with 3×10^7^ FLARE conidia. (B) Representative flow plots of neutrophils from the lungs of B6 and *Mpeg1*^*-/-*^ mice showing RFP (*A. fumigatus* viability) and AF633 (*A. fumigatus* tracer) fluorescence emission. (C) Uptake of conidia by and (D) conidial viability in lung neutrophils, quantified using FLARE conidia and flow cytometry from infected B6 and *Mpeg1*^*-/-*^ mice 24 hpi. (A, C, D) Each dot represents a mouse, and the bar indicates mean. Significance calculated by two-way ANOVA with Šídák’s multiple comparison test. Data are pooled from 2 experiments.

Given that our results conflicted with published data that perforin-2 is essential for murine survival following bacterial infection and for the antibacterial activity of phagocytes (15), we next explored whether findings observed in *Mpeg1*^*-/-*^ mice compared to B6 mice were due to possible differences in the strain background or in the microbiota. To test this possibility, we backcrossed *Mpeg1*^*-/-*^ mice to C57BL/6J mice for 1 generation, then crossed *Mpeg1*^*+/-*^ siblings to generate *Mpeg1*^*-/-*^ and *Mpeg1*^*+/+*^ littermate controls. We first tested the recruitment of phagocytes to the lung and their antifungal activity by infecting *Mpeg1*^*-/-*^ and *Mpeg1*^*+/+*^ controls with FLARE conidia and analyzing lung cells at 24 hpi by flow cytometry. There was no difference in phagocyte numbers in the lung when comparing *Mpeg1*^*-/-*^ mice and *Mpeg1*^*+/+*^ control mice (Fig. 3 A), in contrast to what we observed when comparing *Mpeg1*^*-/-*^ mice to B6 controls (Fig. 2A). Additionally, we did not observe any difference in the uptake or viability of *A. fumigatus* conidia by any population of lung phagocyte between *Mpeg1*^*-/-*^ mice and *Mpeg1*^*+/+*^ control mice (Fig. 3 B-D). In line with our flow cytometry data, we also observed no difference in lung fungal burden by enumerating lung CFUs (Fig. 3 E) and no difference in murine survival (Fig. 3 F) when comparing *Mpeg1*^*-/-*^ mice and *Mpeg1*^*+/+*^ control mice. These findings indicate that perforin-2 is dispensable for phagocyte function during respiratory *A. fumigatus* infection and does not contribute to fungal clearance from the lung. Thus, the observed results that compared *Mpeg1*^*-/-*^ mice to B6 mice were due instead to differences in the strain background or in the microbiota of *Mpeg1*^*-/-*^ mice. Finally, we hypothesized that perforin-2 may be required for clearance of fungal pathogens during a systemic infection, as opposed to mucosal infection. To test this hypothesis, we infected mice with *Candida albicans*, the leading cause of bloodstream fungal infections (1). Mice were infected i.v. with 1.5 × 10^5^ yeast cells and monitored for survival. We found that there was no difference in the survival of *Mpeg1*^*-/-*^ mice compared to *Mpeg1*^*+/+*^ controls (Fig 4). These results suggest that perforin-2 is also dispensable for clearance of *C. albicans* during systemic infections.

**Figure 3.**
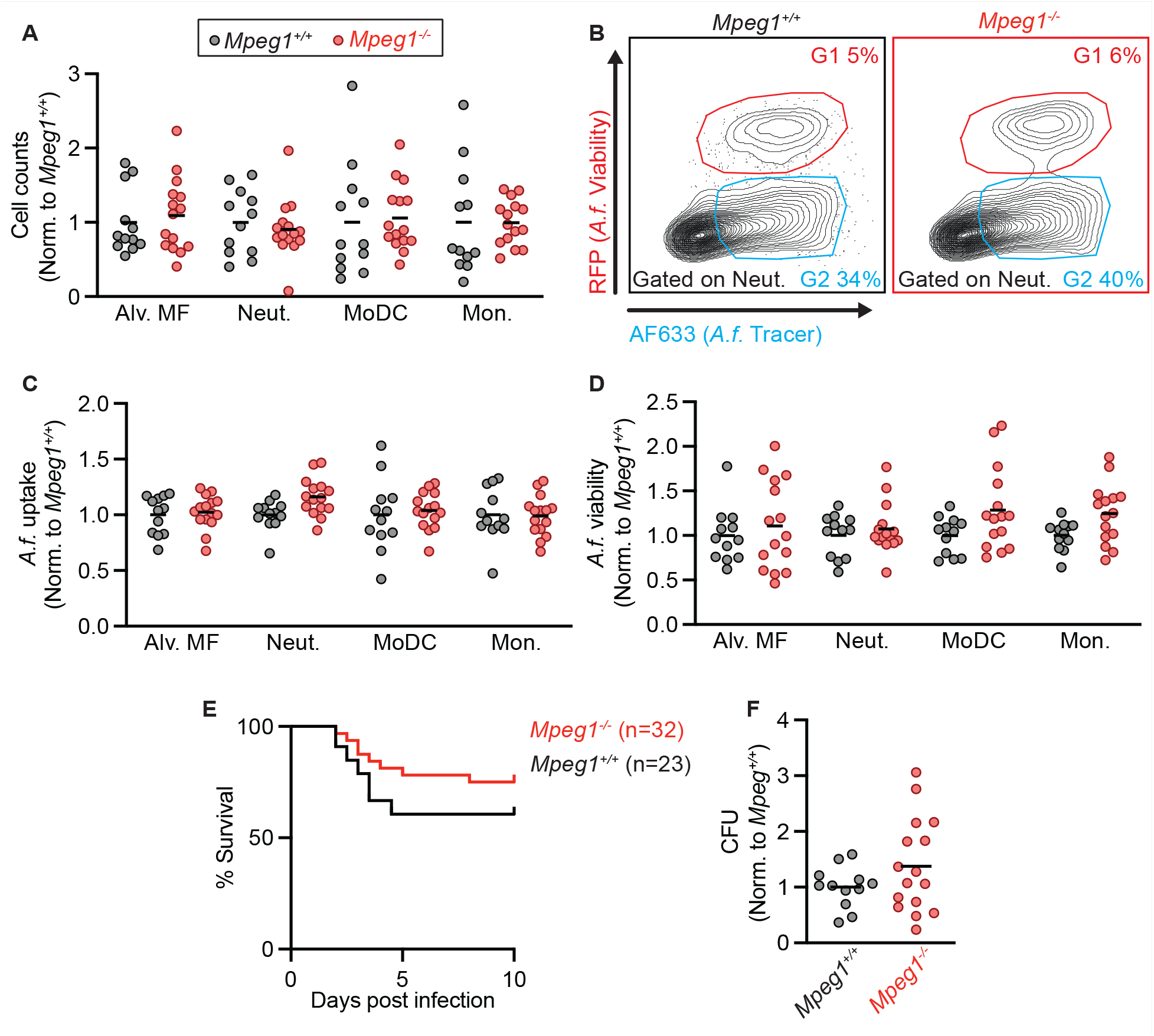
*Compared to littermate controls, phagocytes from* Mpeg1^-/-^ *mice have similar antifungal activity*. (A) Cell counts in the lungs of *Mpeg1*^*+/+*^ and *Mpeg1*^*-/-*^ mice of Alveolar macrophages, neutrophils, MoDCs, and monocytes 24 hpi with 3×10^7^ FLARE conidia. (B) Representative flow plots of neutrophils from the lungs of Mpeg1^+/+^ and Mpeg1^-/-^ mice showing RFP (A.f. viability and AF633 (A.f. tracer). (C) Uptake of conidia by and (D) conidial viability in lung neutrophils, quantified using FLARE conidia and flow cytometry from infected *Mpeg1*^*+/+*^ and *Mpeg1*^*-/-*^ mice 24 hpi. (E) Survival of *Mpeg1*^*+/+*^ and *Mpeg1*^*-/-*^ mice after infection with 4-6×10^7^ *A. fumigatus*. conidia (p=0.186). Significance calculated by log-rank (Mantel-Cox) test. (F) CFU from lungs of *Mpeg1*^*+/+*^ and *Mpeg1*^*-/-*^ mice at 24 hpi with 3×10^7^ conidia. (A, C, D, E) Each dot represents a mouse, and the bar indicates mean. Significance calculated by two-way ANOVA with Šídák’s multiple comparison test. Data are pooled from 2 experiments.

**Figure 4.**
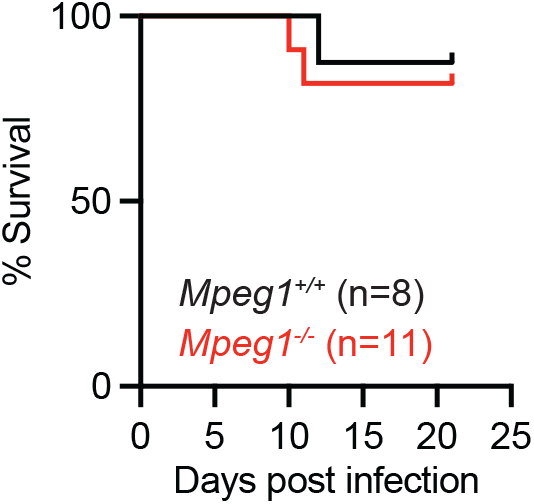
Mpeg1^-/-^ *mice exhibit survival to* Mpeg1^+/+^ *littermates following* C. albicans *infection*. Survival of B6 and Mpeg1^-/-^ mice after i.v. infection with 1.5×10^5^ *C. albicans* yeast cells (p=0.696). Significance calculated by log-rank (Mantel-Cox) test. Data are pooled from 2 experiments.

## DISCUSSION

In this study, we did not uncover an essential role for Perforin-2 in host defense against respiratory *A. fumigatus* and systemic *C. albicans* infections in otherwise immune competent mice. Moreover, perforin-2 is not necessary for phagocyte-mediated killing of *A. fumigatus* conidia. The divergent experimental results observed when comparing *Mpeg1*^*-/-*^ with B6 wild-type or with littermate *Mpeg1*^*+/+*^ control mice indicates that differences the mouse strain background or the microbiota plays an important role in influencing susceptibility to *A. fumigatus* infection.

The role of perforin-2 in antimicrobial immunity remains unclear. While perforin-2 is present in bacteria-containing phagosomes (15, 16) and is associated with the formation of pores on bacterial membranes (15), there are conflicting reports on the susceptibility of *Mpeg1*^*-/-*^ mice to bacterial infection and on its contribution to the bactericidal capacity of phagocytes (15, 16). These contradictory findings on the role of perforin-2 in antibacterial immunity may stem from differences in infection routes and organ-specific immune responses. Our study and Ebrahimnezhaddarzi *et al*. utilized models of intranasal infection with *A. fumigatus* and *M. tuberculosis* or *S. aureus*, respectively, and found no defect in pathogen clearance from the lung (16). In contrast, Mccormack *et al*., which observed a significant effect of perforin-2 deletion on murine survival, employed orogastric and epicutaneous infection models (15).

It is possible that differences in inbred mouse strains may also contribute to different experimental outcomes. Mccormack *et al*. used 129X1/SvJ and mixed C57BL/6J-129X1/SvJ backgrounds (15), while Ebrahimnezhaddarzi *et al*. used a mixed C57BL6/J-BALB/c background for the generation of *Mpeg1*^*-/-*^ mice (16). Despite these differences, both studies used littermate controls, minimizing strain-related variations between experimental and control groups. Our study indicates that *A. fumigatus* infection outcomes may vary due to microbiota or host strain differences. We first used *Mpeg1*^*-/-*^ mice with a mixed C57BL/6J-DBA/2 background (19) prior to generating co-housed littermate controls. Since immunosuppressed DBA/2 mice are more susceptible to *A. fumigatus* than C57BL/6J mice (21), the mixed background of our *Mpeg1*^*-/-*^ mice likely does not explain our observed resistance to *A. fumigatus* compared to C57BL/6J mice.

Because perforin-2 likely inserts into target cell membranes to exert its antimicrobial effect, the susceptibility of different pathogens to perforin-2 may be influenced by the structural composition of their cell walls, which lie outside the cell membrane in fungi and gram-positive bacteria. The cell walls of gram-positive bacteria like *S. aureus* are primarily composed of peptidoglycan, a polymer of alternating N-acetylglucosamine (GlcNAc) and N-acetylmuramic acid (MurNAc) residues, with peptide side chains that cross-link adjacent glycan chains (22). In contrast, the cell walls of *A. fumigatus* conidia consist of a core layer of β-1,3-linked glucan polysaccharides, mannoproteins, galactomannan, and chitin which are further covered by a hydrophobic layer of hydrophobins and melanin under resting conditions (23). These compositional differences in the cell wall may affect the accessibility of perforin-2 to the underlying membrane of target cells, which could explain variations in susceptibility to perforin-2 mediated killing among different pathogens. The thickness of the pathogen cell wall may also contribute to resistance to perforin-2, as fungal cell walls are estimated to be approximately 100 nm thick—at least three times thicker than the cell wall of *S. aureus* (approximately 20-30 nm) (22, 24).

In this study, we found that perforin-2 is dispensable for antifungal immunity during *A. fumigatus* lung infection and *C. albicans* bloodstream infection. These findings expand our understanding of the role of perforin-2 in antimicrobial immunity and contribute to our understanding of which factors are essential and others which are redundant for phagocyte killing of *A. fumigatus* conidia.

## METHODS

### Mice

C57BL/6J mice (stock # 000664) were purchased from The Jackson Laboratory. *Mpeg1*^*-/-*^ have been previously described (19). In brief, *Mpeg1*-deficient mice were created using CRISPR-Cas9 gene editing at the Penn Vet transgenic core facility. Embryos were collected from 6-to 8-week-old superovulated C57BL/6 females, which had been mated with B6D2F1 males (offspring of C57BL/6 females and DBA/2 males). The CRISPR components were microinjected into the cytoplasm of the embryos, which were then transferred to the oviducts of pseudopregnant Swiss Webster female recipients. Mice that were homozygous for the targeted deletion were outcrossed to C57BL/6J mice, and heterozygous offspring were interbred to establish multiple founder lines. The data presented here were derived from a single founder line. All mice used in this study were 8-12 weeks old. Within experiments, mice were age- and sex-matched. Experiments were performed with both male and female mice. Mice were bred and housed in the Research Animal Resource Center at MSKCC in individual ventilated cages under specific pathogen free conditions. C57BL/6J control mice were housed separately from *Mpeg1*^*-/-*^ mice, but *Mpeg1*^*+/+*^ littermates were co-housed with *Mpeg1*^*-/-*^ mice. Animal experiments were conducted with approval of the MSKCC (protocol 13-07-008) Institutional Animal Care and Use Committee. Animal studies complied with all applicable provisions established by the Animal Welfare Act and the Public Health Services Policy on the Humane Care and Use of Laboratory Animals.

### Aspergillus fumigatus strains and murine infection model

*A. fumigatus* strains CEA10 and CEA10-RFP (provided by Robert Cramer, Dartmouth University) were cultured on glucose minimal medium slants at 37°C for 4–7 days prior to harvesting conidia for experimental use. To generate AF633-labeled or FLARE conidia for experimental use, 7×10^8^ CEA10 (for AF633-labeled) or CEA10-RFP (for FLARE) conidia were rotated in 10 μg/mL Sulfo-NHSLC-Biotin (Thermo Scientific) in 1 mL of 50 mM carbonate buffer (pH 8.3) for 2 hours at 4°C, incubated with 20 μg/mL Streptavidin, Alexa Fluor 633 conjugate (Molecular Probes) at 37°C for 1 hour, resuspended in phosphate-buffered saline (PBS) and 0.025% Tween 20 for use within 24 hours. For infections, mice were lightly anesthetized by isoflurane inhalation and 3-6×10^7^ A. fumigatus conidia were instilled via the intratracheal route in 50 μL of PBS + 0.025% Tween-20.

### Quantification of fungal burden

To measure colony-forming units (CFU) in the lungs of infected mice, lungs were dissected and homogenized with a PowerGen 125 homogenizer (Fisher Scientific) for 10-15 seconds in 2 mL of PBS. 10 μL was removed and diluted for plating onto Sabourand dextrose agar plates. Plates were incubated for 48 hours at 37°C and CFU were enumerated by counting.

### Flow cytometry

For analysis of immune cells, single cell suspensions of mouse lungs were generated by putting lungs in gentle MACS C tubes and mechanically homogenizing in 5 ml PBS using a gentle MACS Octo Dissociator (Miltenyi Biotec) in the absence of enzymes, then filtered through 100 μm filters. Next, red blood cells were lysed using RBC lysis buffer (Tonbo Biosciences), cells were blocked with anti-CD16/CD32, stained with fluorophore-conjugated antibodies, and analyzed on a Beckman Coulter Cytoflex LX. Single color controls for compensation were generated using lung cells or OneComp eBeads™ Compensation Beads (ThermoFisher). Experiments were analyzed with FlowJo version 10.8.1. Dead cells were excluded with DAPI or eBioscience™ Fixable Viability Dye eFluor™ 506 (ThermoFisher). Neutrophils were identified as CD45+ CD11b+ Ly6G+ cells, inflammatory monocytes as CD45+ CD11b+ CD11c− Ly6G− Ly6Chi cells, Mo-DCs as CD45+ CD11b+ CD11c+ Ly6G− Ly6Chi MHC class II+ cells, and alveolar macrophages as CD11c+, Siglec-F+. Phagocytes that contain live conidia are RFP+ and AF633+ (G1) and phagocytes that contain dead conidia are RFP-AF633+ (G2). Conidial phagocytosis was quantified as the sum of the fraction of a given phagocyte in the G1 gate and the fraction of a given phagocyte in the G2 gate (G1+G2). To assess how effective phagocytes were at killing conidia, the fraction of viable conidia was calculated as G1/(G1+G2).

### Murine systemic candidiasis infection model

*C. albicans* strain SC5314 was used in this study. Yeast cells were serially passaged 2 times in YPD (yeast extract, bacto-peptone and dextrose) broth, grown at 30°C with shaking for 18-24 hours at each passage. Yeast cells were washed in PBS, counted, and 1.5×10^5^ yeast cells were injected intravenously via the lateral tail vein.

## ACKNOWLEDGEMENTS

We thank all members of the Hohl lab for insightful discussions. We thank Robert Cramer (Dartmouth College) for sharing the *A. fumigatus* strains used in this work. These studies were supported by NIH grants F31 AI161996 (MAA), P30 CA008748 (MSKCC, PI: Selwyn Vickers), R37 AI093808 (TMH), R01 AI139632 (TMH), and U01AI163062 (DRH). The funders had no role in study design, data collection and analysis, decision to publish, or preparation of manuscript.

## Resource Availability

Further information and requests for resources or reagents should be directed to the Lead Contact, Tobias M. Hohl.

